# The effects of ethnoracial-related stressors during pregnancy on the developing offspring brain

**DOI:** 10.1101/2022.08.08.503168

**Authors:** Marisa N. Spann, Kiarra Alleyne, Cristin M. Holland, Antonette Davids, Arline Pierre-Louis, Claire Bang, Victoria Oyeneye, Rebecca Kiflom, Eileen Shea, Bin Cheng, Bradley S. Peterson, Catherine Monk, Dustin Scheinost

## Abstract

We are increasingly aware of the effects of ethnoracial stress on health, with emerging interest in the potential for intergenerational transmission before birth. Here, we investigate the effect of maternal prenatal discrimination and acculturation experiences on fetal growth, birth outcomes, and functional connectivity in the infant brain. In pregnant adolescent women, we collected self-report measures of acculturation (tailored to an adolescent and Latinx population), discrimination, and maternal distress (i.e., stress and depressive symptoms; n=165). Fetal growth were obtained via electronic health records (n=92), and infant amygdala seed connectivity was assessed using functional magnetic resonance imaging (n=38). We found that greater maternal prenatal assimilation to the host culture was associated with slower fetal growth, lower gestational age at birth, and weaker amygdala-fusiform connectivity. Maternal prenatal discrimination was associated with weaker amygdala-prefrontal connectivity. Together, these results suggest intergenerational effects of ethnoracial stressors on the growth and neural development of future generations.

## Introduction

The history of the United States is marked by inequalities amongst people of color(1-3). Yet, recently, the U.S. has been diversifying faster than ever before—the 2010 to 2020 census data reflects an increase in non-white populations (e.g., Latino/Hispanic, Asian and Pacific Islander) and the first decline in the white population in U.S. history(4). Stressors related to ethnoracial inequalities contribute to adverse effects on physical and mental health in populations of color(5). Common ethnoracial stressors include discrimination—the unjust treatment of individuals based on characteristics such as race, gender, age, or sexual orientation(6)—and acculturative stress—the demands, strains, and ‘wear and tear’ of acculturation, or the process of adapting to a new culture(7).

The effects of discrimination and acculturation during pregnancy have profound intergenerational ramifications. Experiences of ethnoracial discrimination are detrimental to the psychological well-being of pregnant women(8) and result in increased infant mortality(9, 10) preterm birth(11), and decreased infant birth weight(12). Similar, experiences of acculturation are associated with depression(13-17), prenatal anxiety(18, 19), and stress—particularly, acculturative stress—in pregnant women(20, 21). Despite the profound impact these experiences have on a pregnant woman of color and their fetuses, there is little research characterizing the impact of these experiences on brain-based outcomes in offspring.

Studies are beginning to demonstrate the neural correlates of ethnoracial stressors and processing in adults(22-24) and the impact of prenatal stressors on infant brain development, which serve as a foundation for understanding the above intergenerational effect. High exposure to discrimination is associated with stronger resting state functional connectivity between the amygdala and several brain regions, including the frontal lobe(25). The amygdala appears to be involved in automatic race evaluations(26). Studies consistently demonstrate that the amygdala is affected by maternal, prenatal distress(27-30). Despite a limited number of existing studies, these results suggest that amygdala connectivity is a good candidate for investigations of brain outcomes associated with prenatal exposure to discrimination and acculturation.

In this secondary data analyses, we investigate the effects of prenatal discrimination and acculturation in pregnant adolescents on fetal growth, birth outcomes, and functional connectivity in the infant brain. To measure maternal experiences during pregnancy, we collected two measures of acculturation (tailored to an adolescent and Latinx population) and an experience of discrimination scale as well as additional measures of maternal distress (e.g., stress and depressive symptoms). We acquired fetal ultrasonographic data to measure fetal growth (i.e., head circumference; n=92), and birth outcome data (i.e., gestational age at birth and Apgar score at 5 minutes; n=155), retrospectively from electronic health records. Using resting-state functional magnetic resonance imaging data acquired at 40–46 weeks post-menstrual age, we assessed amygdala functional connectivity using seed connectivity in a subset of offspring (n=38). We expected maternal experience of discrimination and acculturation will be associated with individual offspring differences in fetal growth outcomes, birth outcomes, and functional brain connectivity.

## Methods and Materials

### Participants

165 nulliparous pregnant women, aged 14 to 19 years, participated in this study. The pregnant women were recruited in the 1^st^ or 2^nd^ trimester through the Departments of Obstetrics and Gynecology at Columbia University Irving Medical Center and Weill Cornell Medical College, and flyers posted in their vicinity as part of a longitudinal study examining adolescent pregnancy behaviors and infant outcomes. The pregnant women received routine prenatal care and had no major health problems at the time of recruitment. Participating women were excluded if they acknowledged the use of recreational drugs, tobacco, alcohol, or medications that affect cardiovascular function, or lacked fluency in English. The women provided informed written consent for themselves and their infant to participate in the study including the infant magnetic resonance imaging (MRI) scan. Infants were imaged within the first 6 weeks of life. All study procedures were approved by the Institutional Review Board of the New York State Psychiatric Institute.

### Self-reported Measures

We collected several measures of acculturation, discrimination, and distress during pregnancy. For acculturation, we collected the Acculturation, Habits, and Interests Multicultural Scale for Adolescents (AHIMSA) and the Short Acculturation Scale for Hispanics (SASH) during the 2^nd^ or 3^rd^ trimester (24–37 weeks of gestation). Acculturation in a population can be represented in one of two ways: a unidimensional model of high versus low assimilation and Berry’s multidimensional model. Berry’s model of acculturation outlines four acculturation categories based on the combination of acquiring/rejecting the host culture and retaining/rejecting the home culture(31). These categories are integration—the identification with both culture, assimilation—the identification with the host culture, separation—the identification with home culture, and marginalization—the identification with neither home nor host culture. The AHIMSA is based on Berry’s model, while the SASH is a unidimensional model. Self-reported language use—a common proxy for acculturation—was also collected. For discrimination, we collected the Experiences of Discrimination (EOD). For distress, we collected the Perceived Stress Scale (PSS) and the Reynold’s Adolescent Depression Scale (RADS). The EOD, PSS, and RADS were collected at three time points during pregnancy: at 12-14, 24–26, and at 34–36 weeks of gestation.

### Electronic Health Record Data

We attained fetal morphometric measures and birth outcomes from participants’ electronic health records. Fetal morphometric measures were head circumference (HC) and biparietal diameter (BDP). HC was measured along the outer perimeter of the calvaria, parallel to the biparietal diameter. BDP was measured from the outer margin of the proximal skull table to the inner margin of the distal skull table. Participants with two or more ultrasonography records were included to construct growth curves of the two primary fetal outcomes. Birth outcomes were gestational age at birth and Apgar score at 5 minutes.

### Imaging Procedures

Using data acquired at 40–46 weeks postmenstrual age, we assessed amygdala functional connectivity using seed connectivity in a subset of offspring. The infants were fed, swaddled, and acclimated to the scanner environment and noise by listening to a tape recording of the scanner sounds played before each pulse sequence. The infants were given time to fall asleep, without the use of sedatives, on the scanner bed before the start of each sequence. Foam and wax earplugs along with ear shields (Natus Medical Inc., San Carlos, CA) were applied to dampen scanner noise. MRI safe ECG and pulse oximetry leads were placed, to monitor heart rate and oxygen saturation throughout the scan (InVivo Research, Orlando, FL). Images were obtained using a 3 Tesla General Electric (GE) Signa MRI scanner (Milwaukee, Wisconsin) and an 8-channel head coil. High resolution anatomical T2-weighted images and functional resting state images were acquired.

#### Common Space Registration

First, anatomical images were skull stripped using FSL (https://fsl.fmrib.ox.ac.uk/fsl/), and any remaining non-brain tissue was removed manually. All further analyses were performed using BioImage Suite unless otherwise specified.

#### Connectivity processing

Motion correction was performed using SPM8 (http://www.fil.ion.ucl.ac.uk/spm/). Images were warped into 3 mm^3^ common space using the non-linear transformation described above and cubic interpolation. Next, images were iteratively smoothed until the smoothness of any image had a full-width half-maximum of approximately 8 mm using AFNI’s 3dBlurToFWHM (http://afni.nimh.nih.gov/afni/). This iterative smoothing reduces motion-related confounds(32). Several covariates of no interest were regressed from the data, including linear and quadratic drifts, mean cerebral-spinal-fluid (CSF) signal, mean white matter signal, and mean gray matter signal. For additional control of possible motion-related confounds, a 24-parameter motion model (including six rigid-body motion parameters, six temporal derivatives, and these terms squared) was regressed from the data. The functional data were temporally smoothed with a Gaussian filter (approximate cutoff frequency=0.12 Hz). A dilated gray matter mask was applied to the data so only voxels within gray matter were used in further calculations. The CSF, white matter, and gray matter masks were manually defined on the reference brain.

#### Seed connectivity

We assessed whole-brain seed connectivity from the right and left amygdala combined into a single seed, shown in Figure S5. Seeds were manually defined on the reference brain. The time course of the reference region in each participant was then computed as the average time course across all voxels in the seed region. This time course was correlated with the time course for every other voxel in gray matter to create a map of r-values, reflecting seed-to-whole-brain connectivity. These r-values were transformed to z-values using Fisher’s transform yielding one map for each seed and representing the strength of correlation with the seed for each participant.

### Statistical Analyses

We examined the factor structure of the four AHIMSA subscales (integration, assimilation, separation, marginalization) and the SASH as a measure of acculturation using the R psych package. Eigen values were first examined to determine the appropriate number of factors, then principal factor analysis was conducted using Pearson’s correlations. Integration and separation subscales were reverse coded to ensure only positive factor loadings. Factor scores were calculated from the final model and used in subsequent main analyses. To investigate the associations between discrimination, acculturation, and distress during pregnancy, we performed an exploratory factor analysis. The scores of scales collected at multiple timepoints were highly correlated (r > 0.5) and, thus, were averaged across trimesters into a single score used in analyses. As the distribution of EOD was highly zero-inflated (52% of women did not endorse discrimination), it was further dichotomized to women experiencing discrimination (EOD>0) and those not. The integration and separation subscales of the AHIMSA were reverse coded to ensure only positive factor loadings.

To assess the associations of the variable of interest with potential confounding variables, demographic and behavioral data were analyzed using standard χ2 test statistics or Fisher exact test for categorical data. Continuous data were analyzed using t tests or Mann-Whitney U tests when a normal distribution could not be assumed to compare groups. Linear regression was used to associate fetal growth and birth outcomes with the ethnoracial stressors. The dependent variables with the three latent factors of ethnoracial stressors (derived from the factor analysis) and discrimination as independent variables were performed. Both unadjusted analyses and analyses adjusted by relevant covariates were performed. All analyses were performed using SPSS statistical software version 25 (IBM) or SAS statistical software version 9.3 (SAS Institute). P values were 2-sided, and P□<□.05 was considered statistically significant.

Imaging data were analyzed using voxel-wise linear models controlling for sex, PMA, and scanner upgrade with all three covariates included in a single model. For primary analysis, the dependent measure was whole-brain functional connectivity of the amygdala. The independent variable were the acculturation factors and experience of discrimination. Secondary analyses were performed to control for maternal depression and perceived stress. Imaging results are thresholded at p < 0.05, with all maps corrected for multiple statistical comparisons across gray matter using cluster-level correction estimated via AFNI’s 3dClustSim (version 16.3.05) with 10,000 iterations, an initial cluster forming threshold of p=0.001, the gray matter mask applied in preprocessing, and a mixed-model spatial autocorrelation function (ACF). Parameters for the spatial ACF were estimated from the residuals of the voxel-wise linear models using 3dFWHMx. The latent factors of ethnoracial stressors (derived from the factor analysis) were used as independent variables while controlling for the neonates age at scan, sex, and scanner upgrade.

## Results

### Demographic Characteristics

The maternal and neonatal demographic characteristics for the full sample (n=165) and the sub-sample with ultrasound data (n=92) and with neonatal MRI data (n=38) are summarized in Table 1. Most pregnant women identified as Hispanic or Latinx (88.48%). Demographic characteristics for the total sample did not differ significantly from the subsample with ultrasonographic data and from the subsample with MRI data.

**Table 1.** Maternal and Neonatal Demographics of Sample with Acculturation Data.

|  |  |  |  |  |  |  |
| --- | --- | --- | --- | --- | --- | --- |
|  | **Factor Analysis**<br>**Sample**<br>**(n=165)** |  | **Sample with**<br>**Ultrasound Data**<br>**(n=92)** |  | **Sample with MRI**<br>**Data**<br>**(n=38)** |  |
| **Variables** | **N** | **Mean (SD)**<br>**or %** | **n** | **Mean (SD)**<br>**or %** | **n** | **Mean (SD)**<br>**or %** |
| *Maternal* |  |  |  |  |  |  |
| Age at delivery, years | 151 | 18.25 (1.35) | 92 | 18.30 (1.39) | 38 | 18.11 (1.47) |
| Pre-pregnancy Body Mass Index (BMI) | 163 | 25.84 (6.96) | 91 | 25.84 (7.29) | 37 | 24.70 (6.47) |
| Years of Education | 163 |  | 90 |  | 37 |  |
| 8th grade | 5 | 3.07% | 1 | 1.11% | 0 | 0.00% |
| 9th grade | 17 | 10.43% | 9 | 10.00% | 5 | 13.51% |
| 10th grade | 20 | 12.27% | 10 | 11.11% | 7 | 18.92% |
| 11th grade | 45 | 27.61% | 26 | 28.89% | 7 | 18.92% |
| 12th grade or higher | 76 | 46.63% | 44 | 48.89% | 18 | 48.65% |
| Ethnicity | 165 |  | 92 |  | 38 |  |
| Not Hispanic/Latina | 19 | 11.52% | 9 | 9.78% | 5 | 13.16% |
| Hispanic/Latina | 146 | 88.48% | 83 | 90.22% | 33 | 86.84% |
| Type of Delivery | 141 |  | 87 |  | 34 |  |
| Vaginal spontaneous | 49 | 34.75% | 37 | 42.53% | 15 | 44.12% |
| Assisted vaginal | 60 | 42.55% | 37 | 42.53% | 12 | 35.29% |
| Emergent C-section | 32 | 22.70% | 13 | 14.94% | 7 | 20.59% |
| Pregnancy Complications | 141 |  | 86 |  | 32 |  |
| None | 102 | 72.34% | 72 | 83.72% | 29 | 90.63% |
| Complications | 39 | 27.66% | 14 | 16.28% | 3 | 9.38% |
| *Neonatal* |  |  |  |  |  |  |
| Gestational Age at Birth, wks | 155 | 39.11 (2.11) | 92 | 39.31 (1.32) | 38 | 39.22 (1.28) |
| Birth Weight, gms | 150 | 3158.09<br>(542.87) | 90 | 3157.35<br>(482.54) | 37 | 3116.42<br>(442.56) |
| Birth Head Circumference, cms | 116 | 33.93 (1.22) | 69 | 33.76 (1.24) | 25 | 33.89 (1.47) |
| Apgar 1 minute | 141 | 8.37 (1.36) | 88 | 8.41 (1.37) | 34 | 8.71 (0.68) |
| Apgar 5 minute | 141 | 8.84 (0.72) | 88 | 8.81 (0.87) | 34 | 8.94 (0.24) |
| Postmenstrual age at scan | --- | --- | --- | --- | 38 | 42.31 (1.56) |
| Gender | 157 |  | 92 |  | 38 |  |
| Female | 65 | 41.40% | 43 | 46.74% | 13 | 34.21% |
| Male | 92 | 58.60% | 49 | 53.26% | 25 | 65.79% |
| All mothers of Hispanic ethnicity were coded together. Abbreviations: cms, centimeters; wks, weeks; gms, grams. |  |  |  |  |  |  |

### Factor Analysis of Ethnoracial Stressors

Our initial factor analysis included the 4 subscales of the AHIMSA (integration, assimilation, separation, and marginalization), the SASH, the EOD, the PSS, and the RADS. Based on the elbow of Scree plot of the eigenvalues, a 4-factor model was determined to be optimal (Fig. S1A). From this analysis, the PSS and RADS scores clustered into a single factor; scores from the AHIMSA and the SASH clustered into three acculturation factors; and EOD did not cluster into a factor (Fig. S1B). Overall, these results suggest that—for the sample at hand— stress and discrimination are single constructs, distinct from acculturation, and that acculturation can be further clustered in multiple factors. Given this, we performed a factor analysis including only the AHIMSA and the SASH. A 3-factor model was determined to be optimal and produced nearly identical factor structure as the previous model (Fig. S1C-D, Fig. S2). From the AHIMSA, low integration and high assimilation clustered into a factor (labeled ASSIMILATION-INTEGRATION). High levels of ASSIMILATION-INTEGRATION reflect a pregnant woman who has come to identify with the host culture (assimilation), rather than integrating both cultures (integration). Separation from the AHIMSA and the SASH clustered into a factor (labeled ASSIMILATION-SEPARATION). High levels of ASSIMILATION-SEPARATION reflect a pregnant woman who has come to identify with the host culture (assimilation), rather than their home culture (separation). Finally, marginalization loaded onto its own factor (labeled MARGINALIZED). Of note, both higher ASSIMILATION-INTEGRATION and higher ASSIMILATION-SEPARATION indicate a higher level of assimilation (i.e., identifying with the host culture). While these factors as similar in the positive direction, they differ in the negative direction. Lower ASSIMILATION-INTEGRATION indicates an integration of both home and host cultures whereas lower ASSIMILATION-SEPARATION indicates higher identification with home culture. Figure 1 places the three observed factors in the context of the four acculturation categories (i.e., assimilation, separation, integration, marginalization). Together, these results suggest that the ethnoracial stressors represent distinct factors from general distress. As such, ASSIMILATION-INTEGRATION, ASSIMILATION-SEPARATION, MARGINALIZED, and EOD were used as primary ethnoracial stressors for remaining analyses.

**Figure 1:**
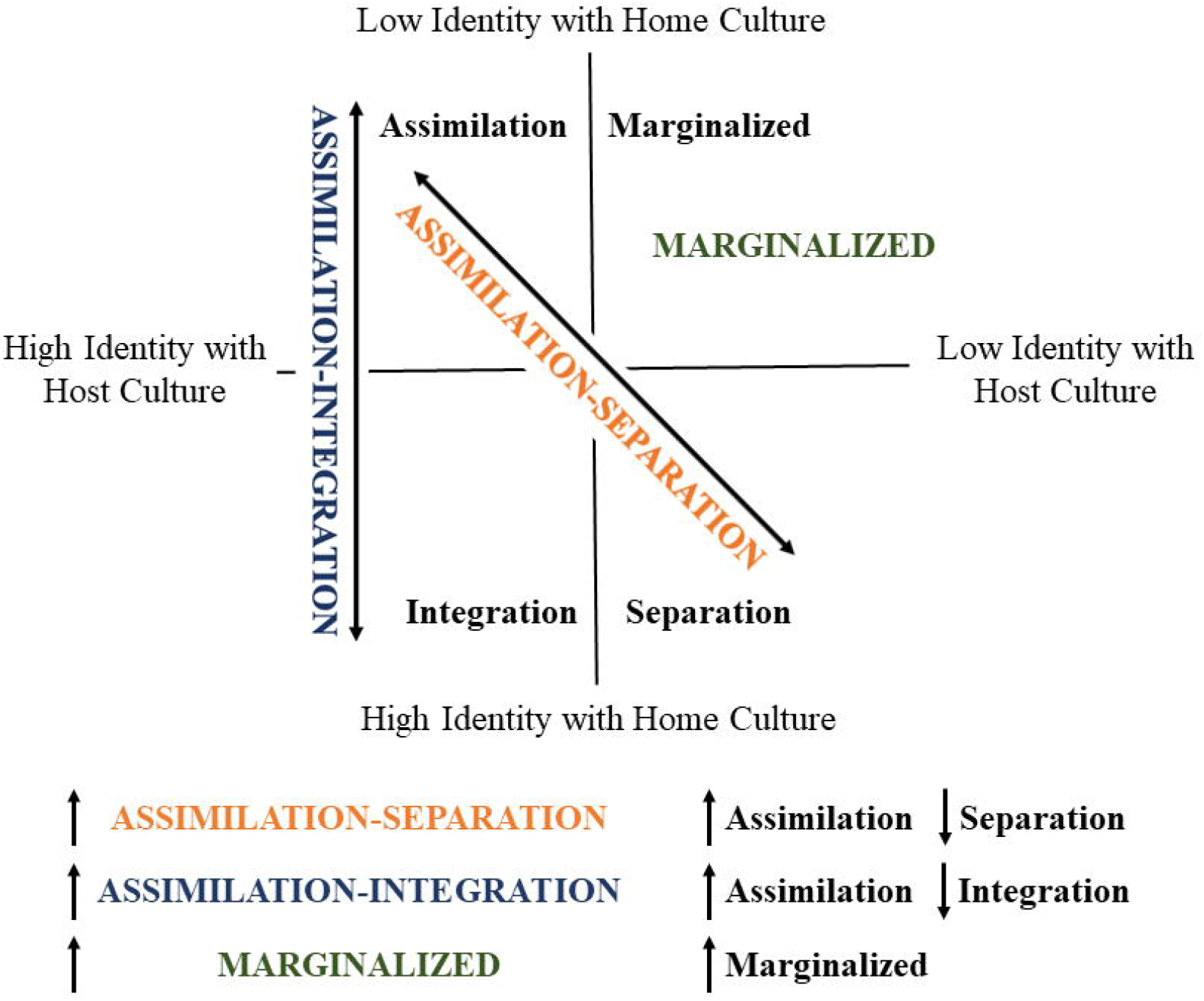
Acculturation factors projected onto the Berry’s model of acculturation. In the Berry’s multidimensional model of acculturation, not all four types may be present in any given population(63), which is reflected in our factor analyses. ASSIMILATION-INTEGRATION and ASSIMILATION-SEPARATION each project onto two categories with the Berry’s model of acculturation. Higher scores in either factor converge to higher assimilation but lower scores on ASSIMILATION-INTEGRATION and ASSIMILATION-SEPARATION diverge to separation and integration, respectively. MARGINALIZED reflected only the marginalized type of acculturation.

While discrimination and maternal distress clustered into different factors, women experiencing discrimination exhibited higher levels of both stress and depressive symptoms (t=1.85, p=0.07, df=163; t=2.23, p=0.03, df=163). There were no significant differences found in acculturation factors between women who experienced discrimination and those who did not experience discrimination during pregnancy (p>0.18).

Language use—a common proxy for acculturation—was associated with the acculturation factors. ASSIMILATION-INTEGRATION was higher in women who did not declare Spanish as either a primary or secondary language compared to women who declared Spanish as their primary language (t= 5.08, p<0.001, df=80) or secondary language (t=-4.7, p<0.001, df=112). In addition, women who declared Spanish as their primary language had lower scores on ASSIMILATION-SEPARATION compared to women who either declared Spanish as their secondary language (t=-8.76, p<0.001, df=132) and women who did not declare Spanish as a primary or secondary language (t=-6.08, p<0.001, df=80).

### Maternal Experiences of Acculturation Associates with Slower Fetal Growth

First, we investigated associations of ethnoracial stressors on fetal growth outcomes. Only higher ASSIMILATION-INTEGRATION was significantly inversely associated with BPD growth (t=-2.06, p=0.04, df=90; Table S1). Models of BDP growth adjusted for maternal demographic variables (maternal age and c-section, ethnicity, and pre-pregnancy body mass index [BMI]) and distress (PSS and RADS) remained significant.

### Maternal Experiences of Acculturation Associates with Poor Birth Outcomes

Next, we investigated associations of ethnoracial stressors on birth outcomes (gestational age at birth and Apgar score at 5 minutes; Table S2). Higher ASSIMILATION-INTEGRATION was significantly inversely associated with Apgar score at 5 minutes (t=-1.98, p=0.049, df=139). Higher ASSIMILATION-SEPARATION was significantly inversely associated with gestational age at birth (t=-2.14, p=0.03, df=153). Models adjusted for maternal demographic variables (maternal age, delivery type, ethnicity, and pre-pregnancy BMI) and distress (PSS and RADS) remained significant.

### Maternal Experiences of Ethnoracial Stressors Associates with Amygdala Connectivity

Finally, we associated the ethnoracial stressors with offspring amygdala connectivity during the neonatal period. As only four women in the imaging sample endorsed experience of marginalization, MARGINALIZED was dropped from this analysis. Higher ASSIMILATION-SEPARATION in mothers was associated with weaker connectivity between the amygdala and bilateral fusiform gyrus (p<0.05; Figure 2A) in their offspring. No significant correlations between for ASSIMILATION-INTEGRATION and amygdala connectivity were observed. In addition, neonates, born to mothers experiencing discrimination, had weaker connectivity between the amygdala and the medial and anterior prefrontal cortices (p<0.05 corrected; Figure 2B). Models adjusted for maternal distress (PSS and RADS) remained significant. Finally, exploratory results using connectivity from the left and right amygdala, independently, exhibited similar results (Figure S3), suggesting that combining the left and right amygdala into a single seed did not influence the results.

**Figure 2:**
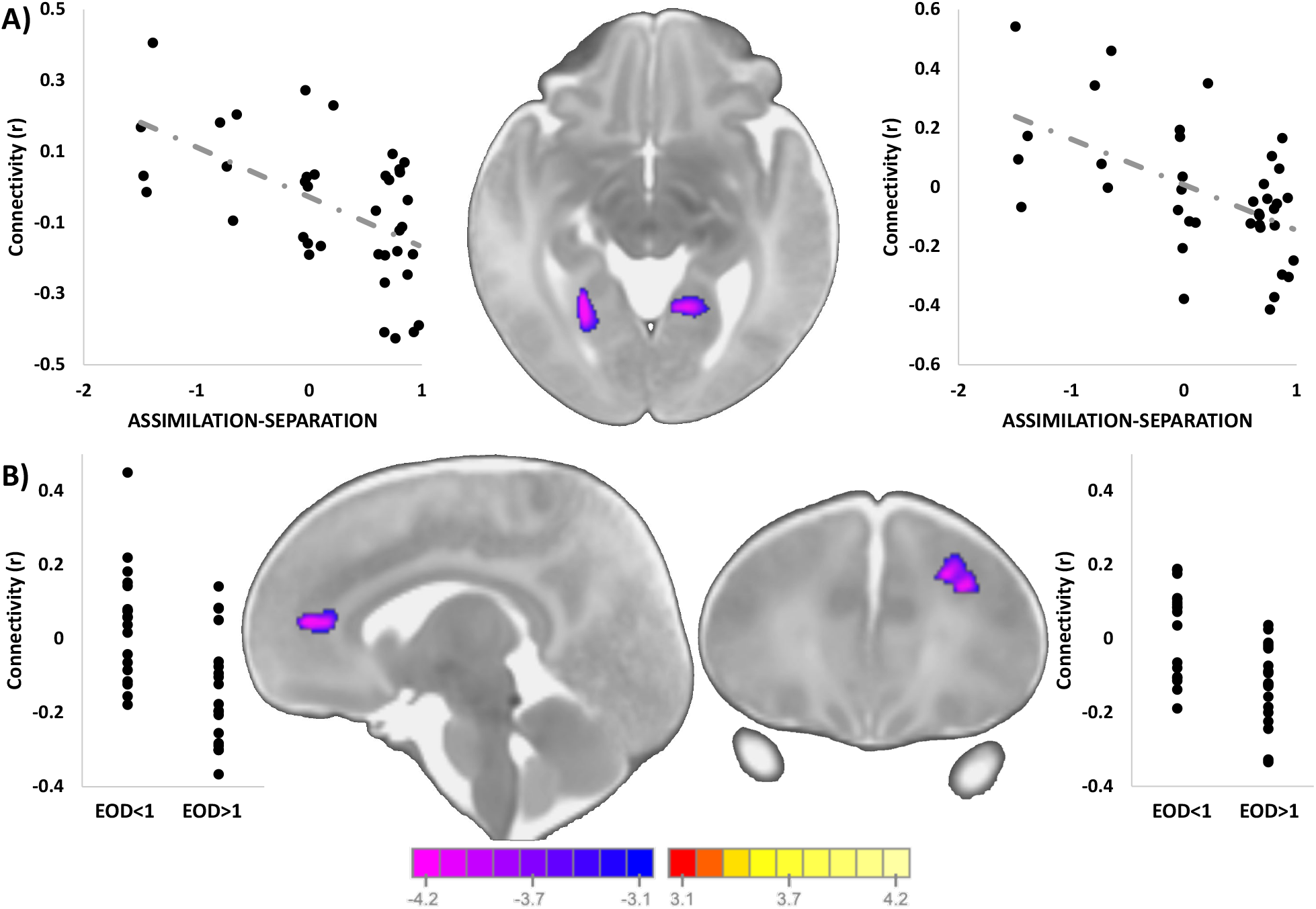
Associations between ethnoracial stressors and amygdala connectivity: **A)** Higher ASSIMILATION-SEPARATION in mothers during pregnancy was associated with weaker infant connectivity between the left and right fusiform cortex. **B)** Neonates, born to mothers experiencing discrimination, had weaker connectivity between the amygdala and the medial and anterior prefrontal cortices. Scatterplots next to the images visualize the distribution of the observed data points for average infant connectivity in the detected regions plotted against ASSIMILATION-SEPARATION or EOD.

## Discussion

This study investigated intergenerational ramifications of ethnoracial stressors during pregnancy. To this end, we examined the effects of acculturation and discrimination experienced by pregnant adolescent women on their offspring’s fetal and birth outcomes, and neonatal amygdala connectivity. Higher assimilation was associated with slower fetal growth, poorer Apgar scores at 5 minutes, lower gestational age at birth, and weaker connectivity between the amygdala and bilateral fusiform gyrus. Maternal experience of discrimination was associated with weaker connectivity between the amygdala and prefrontal cortex. Together, these results demonstrate that the assimilation type of acculturation and experience of discrimination during pregnancy may adversely affect aspects of early growth.

Overall, the associations between assimilation and health outcomes are mixed, leading to a dynamic known as the ‘immigrant paradox’ (21), where recent immigrants, who experience acute acculturative stress, have better health outcomes than immigrants who have been in the United States longer. On the one hand, greater assimilation is associated with increased positive social interactions, which reduces acculturative stress and, putatively, promotes better mental health(33). On the other hand, assimilation has been associated with reduced family support(34) and poorer birth and childhood developmental outcomes(12, 35-37), consistent with our results. As such, greater assimilation is not a simple construct that leads to binary (i.e., good or bad) outcomes and may depend on the sample at hand. Given that our sample is pregnant adolescents and young adults—largely still dependent on their parents or extended family, any reduction in family support could have large-scale effects on their and their child’s health, explaining our results. Thus, it remains important to test these associations in other populations (e.g., older and financially independent, minority women) to fully characterize the effect of acculturation on offspring outcomes.

Functional neuroimaging research has identified the amygdala as being involved in processing in-group and out-membership, as well as race(26, 38, 39). Our finding that greater maternal assimilation is associated with weaker amygdala-fusiform connectivity in offspring is intriguing as previous literature demonstrates an association between the fusiform and race-based processing in adults(40-44). Amygdala-fusiform connectivity also plays a role in ethnoracial processing of faces(39, 41, 45). At three months old, infants are able to start discriminating between own race and other race faces as the “other-race effect” develops(46). As in adults(42), familiarity causes infants to prefer faces of their own race to faces of other races(47), but with greater exposure to other race faces, this preference is negated(48). Further, studies on young infants have shown that the postnatal environmental exposures to people of different ethnoracial groups and cultures affects infants perception of race(49). Our findings may indicate that infants with mothers who report increased identification with their home culture are mainly exposed to faces of their own race during early postnatal life, while greater maternal assimilation to the host culture may provide the infant greater exposure to other race faces. This early life exposure to other race faces may induce changes to brain function related to race processing (such as differences in amygdala-fusiform connectivity) before external behaviors are observable. Nevertheless, follow-up data during infancy would be needed to test these potential links.

Altered amygdala-frontal circuitry is a common sequelae of early life adversity from the pre- and post-natal environment(50-56). Our results are consistent with previous reports of prenatal and other life stressors altering amygdala-frontal circuitry(57) and add maternal experiences of discrimination during pregnancy as an additional stressor that may affect this circuit. Of note, the association between experiences of discrimination and amygdala-frontal circuitry appears to be independent of maternal stress and depressive symptoms, both of which are known to affect this circuitry(50-56). This suggests specificity to experiences of discrimination. As there are many overlapping factors that may alter amygdala-frontal connectivity, understanding how these experiences have shared or unique effects is needed. For example, in our sample, women who experienced discrimination exhibited slightly greater stress and depression. However, the causal nature of these associations is not understood (e.g., does discrimination lead to higher depressive symptoms?). The experience of discrimination may also lead to other feelings, such as anger. While this prenatal exposure is not as well researched as stress and depression in studies that include infant neuroimaging(57), high exposure to prenatal maternal anger has been shown to negatively affect offspring(58) and thus measures related to anger could be explored in future studies. Overall, understanding the effect of maternal experiences of discrimination on the developing brain unique from other negative maternal emotions remains a critical next step.

Experiences of discrimination did not cluster with acculturation. An important consideration about our measure of discrimination is that it is based on self-perception. A person’s affirmation that they have been discriminated against is dependent not only on the occurrence of a negative action based in prejudice but the perception of the action as such. Any variations in perception could lead to under- or over-reporting. While there was not a significant difference in acculturation between those who experienced discrimination and those who did not, prior work indicates that acculturation could have a hand in moderating the experience of discrimination, such that as an individual becomes more integrated into their host society their perception of discrimination becomes heightened(59, 60).

There were several key strengths in our study. MRI scans were acquired within the first few weeks of birth; thus, our findings are largely attributable to the prenatal environment rather than the postnatal environment. Our sample consisted of predominately Latinx participants, allowing for the collection of data from a population that may experience discrimination and acculturation and is underrepresented in neuroimaging and biomedical studies. Our unique approach to measuring acculturation consisted of a combination of two scales, the SASH and AHISMA, measures that have good construct validity(61, 62) and go beyond prior studies’ utilization of language as a primary determinant of acculturation. Furthermore, our study focuses on discrimination as a separate entity from stress, an important distinction to be made given our sample and the current social climate.

Our study also had several limitations. Firstly, our maternal sample consisted of predominantly adolescent pregnant women most of whom were of Latinx descent but excluded participants who lacked fluency in English which limits the generalizability of the findings. Further, acculturation studies have limits with respect to generalizability given that every culture is different and people within each ethnoracial group are not homogonous. Most of our infant sample were male; however, the design of our study was underpowered to detect sex differences. Finally, our study utilized secondary data analysis. Future studies should employ a prospective study design, as well as a design that accounts for measures reflecting acculturative stress, microaggressions, stigma, and family support to better understand acculturation pathways as well as follow-up ethnoracial processing data in the infants.

As our society strives for greater cultural inclusivity and sensitivity, we must become aware of the effects that the experience of acculturation and discrimination have on pregnant women and their infants. Our findings suggest that maternal prenatal discrimination and acculturation are additional stressors that negatively affect fetal growth and birth outcomes, while also associating with the neonatal functional connectivity of the amygdala. Going forward, studies would benefit from the inclusion of these ethnoracial stressors to further our understanding of the intergenerational effect of these stressors on the growth and neural development of future generations.

## Acknowledgements

Data were provided by the National Institute of Mental Health (NIMH) MH093677 (C.M. & B.S.P.) grant. This work was also supported by NIMH K24MH127381 (M.S.), R01MH117983 Supplements 1 (A.PL.) and 2 (A.D.); the National Center for Advancing Translational Sciences TL1TR001875 (C.H.); National Health and Lung and Blood Disease Institute R25HL096260, the BEST-DP: Biostatistics & Epidemiology Summer Training Diversity Program (R.K. & V.O.); Eunice Kennedy Shriver National Institute for Child Health and Human Development K23HD092589 (M.S.); and the Irving Institute for Clinical and Translational Research at Columbia University Irving Medical Center, Irving Scholar Award (M.S.). We also wish to thank Kristiana Barbato, Dr. Ezra Aydin, and Dr. Angeliki Pollatou for helpful feedback on the manuscript concepts, and Dr. Seonjoo Lee for providing initial consultation regarding the statistical analysis.

