## Supplemental Methods, Figures, and Tables for "The effects of ethnoracial-related stressors during pregnancy on the developing offspring brain"

**SI Methods**

***Demographics*.** A previously validated psychosocial questionnaire was administered to each participant to obtain demographic information such as ethnicity, education, and family structure. Prenatal electronic health records were reviewed to determine birth outcomes including gestational age at birth, birth weight, infant length, Apgar score, and delivery data. Obstetrical and neonatal records were reviewed to extract information on the postmenstrual age (PMA), Apgar scores, gestational age at birth, birth weight, and delivery data from medical records. PMA is defined as the time elapsed between the first day of the pregnant woman’s last normal menstrual period and the time of the MRI scan of their infant. Apgar is an assessment performed on the neonate following birth at 1- and 5-minutes following birth. The score includes breathing effort, heart rate, muscle tone, reflexes, and skin color. The total score ranges from 1 to 10. Gestational age at birth was determined from the medical record for dates of ultrasound examinations and last reported menstrual cycle.

***Acculturation.*** Acculturation can include any individual changes that occur due to migration and subsequent interactions with differing cultures^64,65^. Level of acculturation was obtained with two measures—the Short Acculturation Scale for Hispanics (SASH) and the Acculturation, Habits, and Interests Multicultural Scale for Adolescents (AHIMSA).

The SASH measures behavioral factors previously associated with acculturation, including language use, media preference, and ethnic social relations^66^. This measure has 3 subscales and 12-items. The language use subscale asks participants which language they use when reading, speaking, with a child, at home, thinking, and with friends. The media use subscale asks participants to identify the language with which they watch and listen to TV programs, radio programs and movies, and the ethnic social relations subscale asks participants who they spend time with and relate to most. Total scores range from 12-60 with higher scores indicating greater levels of acculturation. SASH has been psychometrically examined and has been found to be both a useful and valid measure of acculturation^62^.

The AHIMSA is a measure of acculturation^63^ for adolescents. It has 8 questions that address adolescents’ social experiences, such as who they are friends with and where their TV shows come from. There are four response categories which each indicate a varying level of acculturation: a. The United States (Assimilation), b. The country my family is from (Separation), c. Both (Integration), d. Neither (Marginalization). Four scores are given based on the four response categories or orientations, and the score for each orientation ranges from 0-8.

***Discrimination***. The Experience of Discrimination (EOD) instrument measured self-reported experiences of ethnoracial discrimination and was scored by counting the number of situations for which a participant experienced ethnoracial discrimination^67^. The validity and reliability of the measure has been tested among a diverse population in order to confirm its psychometric properties^68^. As the distribution of EOD was highly zero-inflated (52% of women did not endorse discrimination), we dichotomous the scale into women not experiencing discrimination (EOD=0) and women experiencing discrimination (EOD>0).

***Perceived Stress.*** The Perceived Stress Scale (PSS) is a commonly used measure for subjective stress with good internal and test-retest reliability^69^. It measures perceived stress by having participants rate their feelings related to potentially stressful events that had occurred within the last month. There are 14 items in which 7 of the items are negative and 7 of the items are positive. Each item was rated on a five-point Likert-type scale in which 0 indicated never and 4 indicated very often. The 7 positive items were then reverse coded so that all the scores could be summed. Higher scores demonstrated greater stress.

***Depression.*** The Reynolds Adolescent Depression Scale (RADS) is a self-report measure that reflects symptoms of depressive disorders in adolescents^70^. The 30–item scale has four subscales: Dysphoric Mood, Anhedonia/Negative Affect, Negative Self-Evaluation, and Somatic Complaints. Item scores from each subscale are summed to create a composite depression score that ranges from 30-120, higher scores are an indication of more severe depressive symptoms. RADS is a commonly used measure due to its proven validity and reliability^71^.

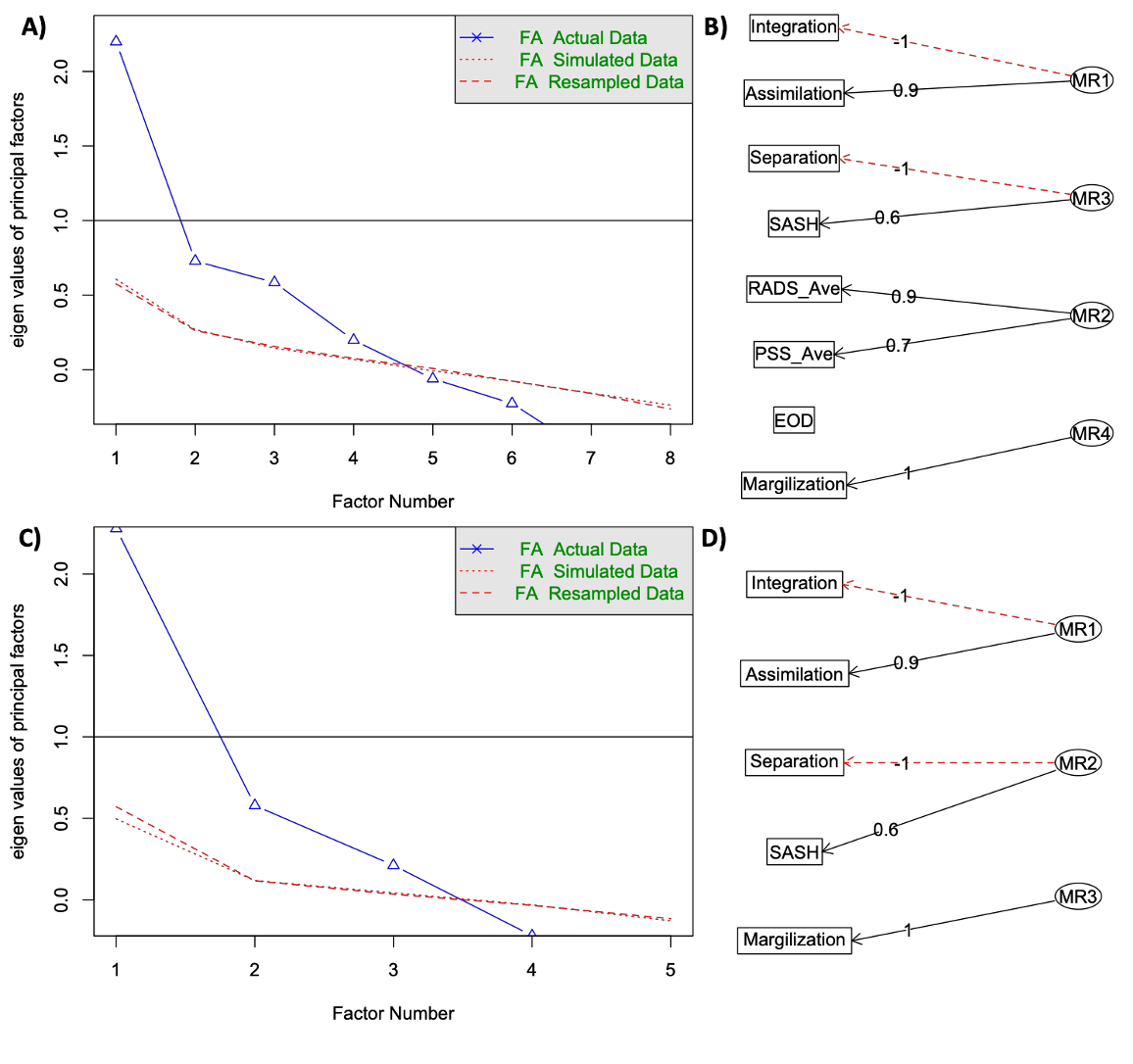

**Figure S1: Factor analysis results. A)** For the factor analysis of experienced distress, acculturation, and discrimination during pregnancy, four factors were determined to be the optimal solution. **B)** Stress clustered into a single factor, while acculturation clustered into three factors. Discrimination did not cluster into a factor. **C)** For the factor analysis of experienced acculturation during pregnancy, three factors were determined to be the optimal solution. **D)** Similar acculturation factors were found with this reduced model compared to the full model that included distress and discrimination. Solid black lines in **B)** and **C)** indicate a positive loading of a measure on a factor, while red dot lines indicate a negative loading.

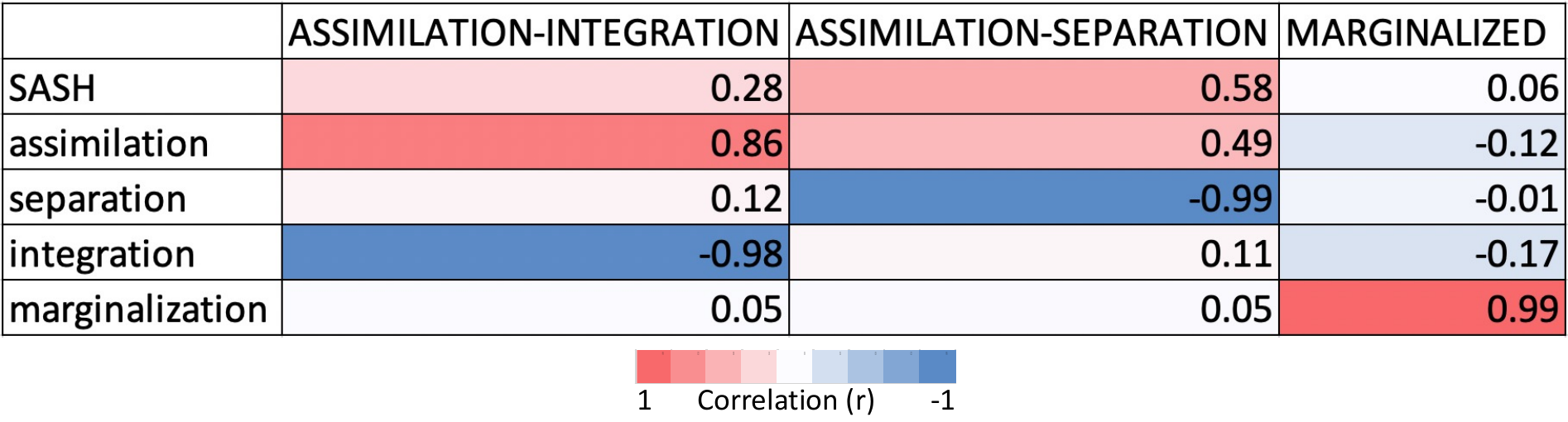

**Figure S2: Correlation matrix between the acculturation measures and acculturation factors derived from the factor analysis.** As suggested by the factor analysis, ASSIMILATION-INTEGRATION was strongly positively correlated with assimilation and inversely correlated with integration. ASSIMILATION-SEPARATION was strongly positively correlated with SASH and assimilation and inversely correlated with separation. Finally, MARGINALIZED was only positively correlated with marginalization.

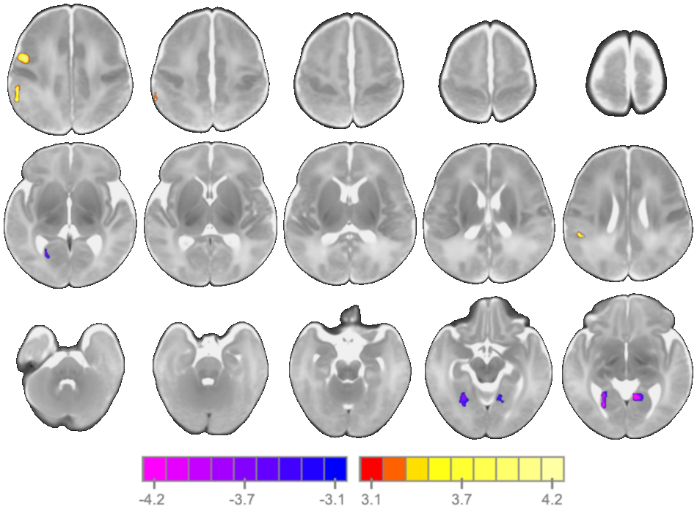

**Figure S3: Associations between ASSIMILATION-SEPARATION and amygdala connectivity for only the left amygdala seed.** Similar results were observed with the left amygdala seed, suggesting that combining the left and right amygdala into a single seed did not influence our results.

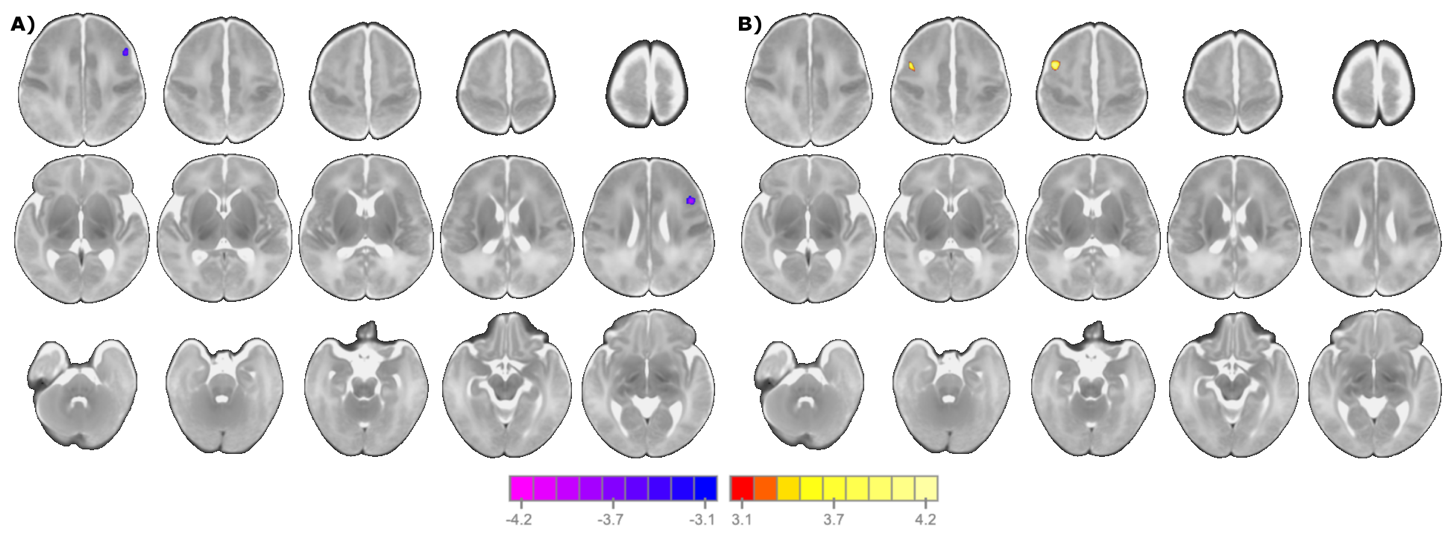

**Figure S4: Associations between EOD and amygdala connectivity for the A) left and B) right amygdala seeds, independently.** Similar results were observed with the left amygdala seed, suggesting that combining the left and right amygdala into a single seed did not influence our results.

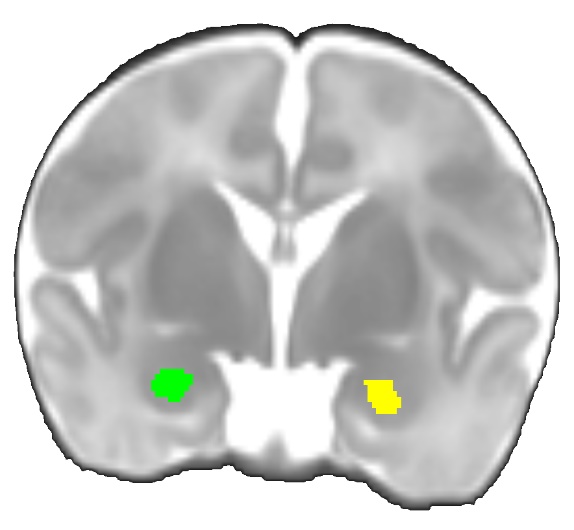

**Figure S5: Left (shown in green) and right (shown in yellow) amygdala seeds.** For the main analyses, these seeds were combined into a single seed for whole-brain connectivity

| **Table S1: Associations between ethnoracial stressors and fetal growth outcomes** | | | | |
| --- | --- | --- | --- | --- |
| **Ethnoracial Stressors** | **Fetal Growth Outcomes** | | | |
|  | **Head Circumference** | | **Bi-Parietal**  **Diameter** | |
|  | ***T*** | **p-value** | ***t*** | **p-value** |
| ASSIMILATION-INTEGRATION | 0.50 | 0.62 | **-2.06** | **0.04** |
| ASSIMILATION-SEPARATION | 0.67 | 0.50 | 1.30 | 0.20 |
| MARGINALIZED | -0.99 | 0.33 | -0.84 | 0.40 |
| Experience of Discrimination | 0.75 | 0.46 | 1.15 | 0.25 |

| **Table S2. Associations between ethnoracial stressors and birth outcomes** | | | | |
| --- | --- | --- | --- | --- |
| **Ethnoracial Stressors** | **Birth Outcomes** | | | |
|  | **Gestational Age at Birth** | | **Apgar Score at**  **5 minutes** | |
|  | ***T*** | **p-value** | ***t*** | **p-value** |
| ASSIMILATION-INTEGRATION | -0.04 | 0.97 | **-1.98** | **0.05** |
| ASSIMILATION-SEPARATION | **-2.14** | **0.03** | -1.48 | 0.12 |
| MARGINALIZED | 0.15 | 0.88 | -0.70 | 0.49 |
| Experience of Discrimination | -1.53 | 0.13 | -0.77 | 0.44 |
